# Assessing the Suitability of Deubiquitylases As Substrates For Targeted Protein Degradation

**DOI:** 10.1101/2025.06.13.659525

**Authors:** Joel Tong, J. Monty Watkins, James M. Burke, Thomas Kodadek

## Abstract

The development of selective inhibitors of Deubiquitylase enzymes (DUBs) is difficult due to a high level of homology in the active sites of the ≈ 100 such enzymes in the human proteome. A potential way to achieve this in a more facile manner would be to develop PROTACs or molecular glues that engage the target DUB in a less conserved region outside of the catalytic domain. However, this raises the concern that auto-deubiquitylation would make DUBs poor substrates for this modality. Here we describe a chemical genetics system to evaluate this issue. We find that some DUBs are readily degradable via the Ubiquitin-proteasome pathway and some are not. Of the latter category, some resist turnover through auto-deubiquitylation and some are simply poor proteasome substrates.

**Significance:** Deubiquitylases (DUBs) are a family of specialized proteases that hydrolyze the isopeptide bond between a lysine and the C-terminal carboxylate of Ubiquitin. DUBs are involved in a myriad of cellular processes and many are attractive drug targets. However, it is difficult to develop selective orthosteric inhibitors due to the high degree of homology between DUB active sites. Targeted protein degradation using a proteolysis-targeting chimera (PROTAC) that recognizes the DUB in a less conserved region outside of the catalytic domain constitutes an attractive alternative strategy for selectively inhibiting a given DUB. Such ligands may not block the catalytic activity of the enzyme, raising the concern that auto-deubiquitylation will make DUBs inherently poor substrates for PROTACs of this type. Since drug-like ligands that engage DUBs outside of the active site are extremely rare, this issue is difficult to address in a straightforward fashion. In this study we establish a generally applicable chemical genetics workflow to evaluate the degradability of DUBs by a PROTAC. The data indicate that some DUBs are readily degradable and some are not. In particular, USP11, an attractive drug target in various cancers and Alzheimer’s disease, is shown to be rapidly degradable, while its paralogs, USP4 and USP15 resist degradation through auto-deubiquitylation.

## Introduction

Ubiquitylation is a key post-translational modification (PTM) that has diverse functional outcomes, the most well-known being K48-linked ubiquitylation that targets a protein for proteasomal degradation^1^. This is a reversible PTM. Ubiquitylation is mediated by E1, E2 and E3 ubiquitin ligases, which create an isopeptide bond between a lysine side chain and the C-terminal carboxylate of Ubiquitin, while deubiquitylases (DUBs) are proteases that cleave these isopeptide bonds^2^. There are almost 100 DUBs encoded in the human proteome, and many of these are critical players in disease progression. For example, USP7 reduces the stability of the tumor suppressor p53 by deubiquitylating its E3 ligase MDM2^3,4^. USP11 has been shown to promote tumorigenesis by deubiquitylating and hence stabilizing oncogene products such as c-Myc and PGAM5^5,6^. Beyond oncology, USP30 has been linked to Parkinson’s disease due to its role in counteracting Parkin-mediated mitophagy^7^, while inhibition of USP14 was demonstrated to enhance turnover of the aggregation-prone Tau protein that is relevant in Alzheimer’s disease^8^. Hence, many DUBs represent attractive drug targets.

As most DUBs are cysteine proteases, it is relatively straightforward to develop a small molecule inhibitor targeting the active site. However, achieving selectivity, particularly across the large USP family, is a major challenge as these DUBs share a high degree of homology in their catalytic domain. This obstacle has been overcome in a few cases, for example the development of highly selective, active site-targeted inhibitors of USP7, but these were the result of major medicinal chemistry efforts^9–11^. Recently, selective inhibitors for USP9X^12^ and USP30^13^ have been reported, but these were also highly labor- and time-intensive endeavors. There is therefore a pressing need for more facile development of DUB therapeutics.

Buhrlarge and co-workers reported an interesting chemoproteomics platform to address this selectivity challenge^14^. Compounds in their library had an electrophilic warhead to target the catalytic cysteine and various heterocyclic building blocks to target the DUB at a less conserved surface some distance away from the active site. Different linkers were used to connect the two. Through this work, they identified a selective inhibitor of VCPIP1 with an IC_50_ of 70 nM. However, it is notable that most of the compounds screened had poor selectivity, especially across the USP family.

An alternative approach that would avoid this daunting selectivity problem would be to employ a targeted protein degradation strategy using a proteolysis-targeting chimera (PROTAC) that engages the DUB of interest in a less conserved domain away from the catalytic site. For example, USP11 has a DUSP domain that is present in only seven out of the approximately 100 DUBs. However, it is reasonable to anticipate ligands that engage a DUB away from the active site will not inhibit its catalytic activity. This, in turn, raises the obvious concern that DUBs will be poor substrates for this type of PROTAC due to auto-deubiquitylation, which is a known phenomenon for some DUBs^15–19^. Unfortunately, a straightforward assessment of this issue is complicated by the fact that there are almost no DUB-binding small molecules that engage the protein outside of the active site. Therefore, we sought to develop a chemical genetics-based workflow to address this issue and “de-risk” PROTAC development campaigns against a DUB of interest. In this study, we use the dTAG system^20^ to investigate the degradability of catalytically active DUBs and specifically assess the degree to which auto-deubiquitylation acts to shield these enzymes from targeted protein degradation. We show that a FKBP12^F36V^-USP11 fusion protein is highly degradable by the PROTAC dTAG-13, while FKBP12^F36V^ fusions of USP4, USP15 and UCHL1 are only modestly degraded. For USP4 and USP15, that this resistance to degradation is largely due to auto-deubiquitylation, whereas UCHL1 appears to simply be a poor substrate for proteasome-mediated proteolysis due to its compact structure. Based on these results, we propose three main categories for DUBs with respect to their suitability as PROTAC substrates and discuss the possibility of exploiting auto-deubiquitylation to selectively degrade USP11 over its paralogs USP4 and USP15.

## Results

### FKBP*-USP11 is readily degradable by the PROTAC dTAG-13

To interrogate the degradable DUB-ome, one approach would be to utilize a pan-DUB PROTAC that recruit DUBs to an E3 ubiquitin ligase and assess degradation via a chemoproteomic workflow. This strategy has previously been adopted to probe the degradability of various kinases and SLC transporters^21–24^. However, all currently available pan-DUB ligands are active site-targeted inhibitors^25,26^, which would preclude the assessment of DUB degradability when the DUB is not inhibited. In addition, as mentioned above, there are almost no selective, non-inhibiting DUB ligands that would allow us to test each DUB individually.

Due to the dearth of suitable tool compounds, we adopted the dTAG technology for our investigation. Briefly, a mutant FKBP tag (FKBP12^F36V^), henceforth referred to as FKBP*, is fused to a protein of interest (POI). The chemical dimerizer dTAG-13 binds to FKBP* and the E3 ligase Cereblon, inducing proximity between the two. This leads to the poly-ubiquitylation and subsequent proteasomal degradation of the POI^20^. In this way, the POI is targeted for degradation without disrupting its catalytic activity. FKBP* was fused to the DUBs USP11, USP4, USP15 and UCHL1. An FKBP fusion with HDAC1 was also constructed. All of the constructs included a C-terminal FLAG tag (Figures 1A and 1B). The DUBs USP11, USP4, USP15 and UCHL1 were chosen because they are known to play important roles in cancer^5,6,27–32^. HDAC1 was chosen as a control because it is an amide hydrolase, but not a DUB, and is readily degradable using the dTAG system^20^. These FKBP- and FLAG-tagged fusion proteins were stably expressed in Flp-In 293 cells. Upon treatment of these cells with dTAG-13, degradation of the FKBP*-DUB-FLAG constructs was evaluated by Western blotting using anti-FLAG antibody.

**Figure 1.**
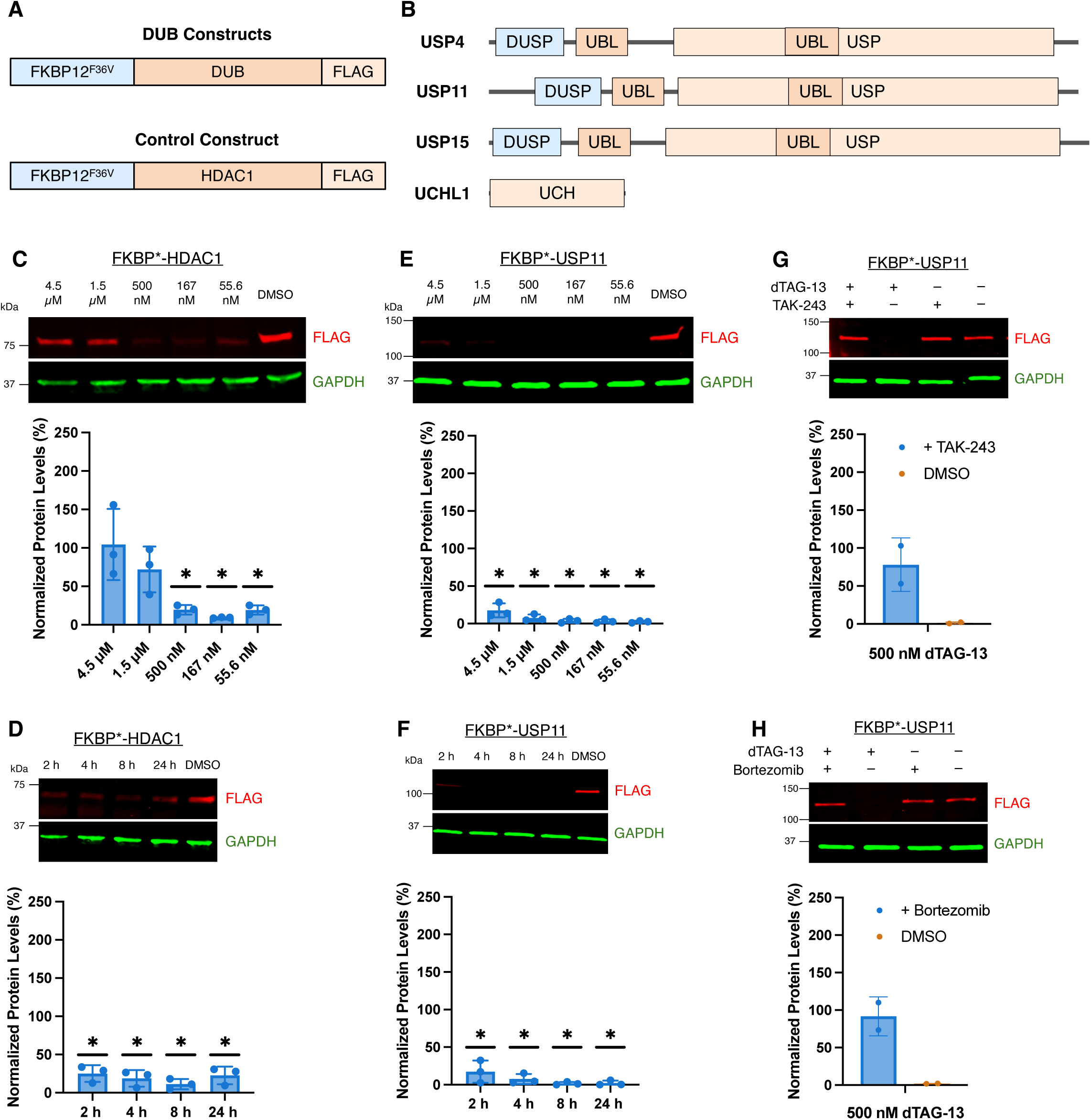
FKBP*-USP11 is readily degradable by the PROTAC dTAG-13. (A) General structure of the FKBP*-tagged proteins that were tested in this study. (B) Schematic of the domain architecture of the DUBs that were tested in this study. (C and E) Representative western blot and quantification of FKBP*-HDAC1 (C) or FKBP*-USP11 (E) levels after HEK293 cells stably expressing the indicated FKBP* construct were treated for 4 h with a range of dTAG-13 concentrations (55.6 nM – 4.5 µM). Data are presented as mean ± SD (n = 3 biological replicates), unpaired t test with vehicle-treated cells (*q < 0.01). (D and F) Representative western blot and quantification of FKBP*-HDAC1 (D) or FKBP*-USP11 (F) levels after HEK293 cells stably expressing the indicated FKBP* construct were treated with 500 nM dTAG-13 for 2 h, 4 h, 8 h or 24 h. Data are presented as mean ± SD (n = 3 biological replicates), unpaired t test with vehicle-treated cells (*q < 0.01). (G and H) Representative western blot and quantification of FKBP*-USP11 levels after HEK293 cells stably expressing the indicated FKBP* construct were treated with 500 nM dTAG-13 ± 10 µM TAK-243 (G) or 10 µM Bortezomib (H) for 4 h. Data are presented as mean ± SD (n = 2 biological replicates).

As expected, there was significant degradation of the control construct FKBP*-HDAC1 at an optimal concentration of dTAG-13 (167 nM; Figure 1C). At higher dTAG-13 concentrations, degradation was much less efficient due to the hook effect, where the PROTAC primarily forms unproductive binary complexes instead of a ternary complex.^33^ FKBP*-HDAC1 degradation by dTAG-13 was also fast, with depletion to 25 % of DMSO levels observed within two hours (Figure 1D).

The FKBP*-USP11 fusion protein was also degraded efficiently. Across all the concentrations of dTAG-13 tested, protein levels were less than 25 % of that in vehicle-treated cells after four hours, reaching as low as 2.5 % with 55.6 nM dTAG-13 (Figure 1E). This degradation was rapid. DUB levels were about 17 % relative to those seen in vehicle-treated cells after just two hours of dTAG-13 treatment (Figure 1F). Co-addition of the E1 inhibitor TAK-243 (Figure 1G) or the proteasome inhibitor Bortezomib (Figure 1H) with dTAG-13 abolished FKBP*-USP11 turnover. These data indicate that USP11, at least in the context of this system, is readily degradable by a PROTAC via the Ubiquitin-proteasome pathway.

### FKBP*-USP11 is active

The fact that FKBP*-USP11 was so readily degradable is significant as it is a high priority cancer target. However, a concern is that the fusion protein has reduced catalytic activity, allowing it to be degraded more readily than would be the case for native USP11. The deubiquitylase activity of USP11 plays an important role in DNA double-strand break (DSB) repair^34,35^. Cortez and co-workers have shown that knocking down USP11 alone, even in the absence of ionizing radiation, was sufficient to cause a three- to four-fold increase in γγ-H2AX foci^34^, which is a marker of unrepaired DSBs^36^. We thus assessed if FKBP*-USP11 complements the loss of USP11 in DSB repair after knockdown of the endogenous protein.

Endogenous USP11 was knocked down using siRNA targeting the untranslated region of USP11 (UTR1 siRNA). In one set of cells, FKBP*-USP11 was also depleted by treatment with dTAG-13, while in a parallel set of cells FKBP*-USP11 levels were left unperturbed. If FKBP*-USP11 is able to function in DSB repair, we would expect to see a higher level of γ-H2AX foci in the cells where both proteins were knocked down than in cells where only endogenous USP11 was depleted. We confirmed by Western Blot that endogenous USP11 species was depleted in each sample (Figures 2A and 2B). We found that there was a marked increase in γ-H2AX foci when both endogenous USP11 and FKBP*-USP11 levels were depleted, as expected. In addition, the more compact nuclei in these cells could be attributed to chromatin condensation, which is an integral component of the DNA damage response^37^. However, in cells where only endogenous USP11, but not FKBP*-USP11, was depleted, this effect was attenuated (Figures 2C and 2D, Figures S1A and S1B). These data show that FKBP*-USP11 can buffer the deleterious loss of endogenous USP11, suggesting that it is a functional protein with respect to DSB repair.

**Figure 2.**
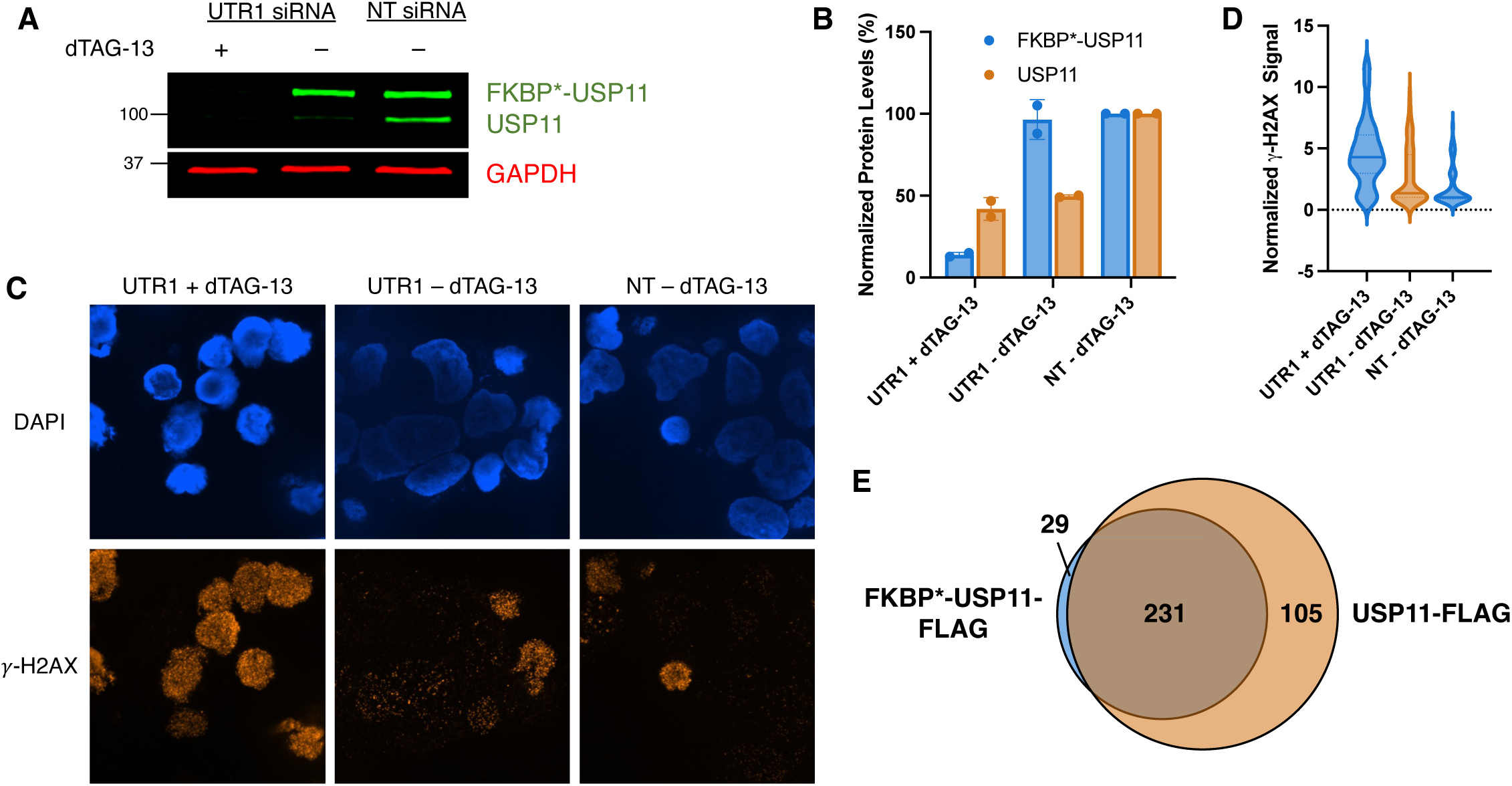
FKBP*-USP11 is active. (A) Western blot of FKBP*-USP11 and endogenous USP11 levels in HEK293 cells stably expressing FKBP*-USP11 after the indicated treatment. Cells were transfected with siRNA targeting the untranslated region of USP11 (UTR1 siRNA) or non-targeting siRNA (NT siRNA) (72 h incubation post-transfection) and treated with 500 nM dTAG-13 or vehicle control (24 h treatment). Blot is representative of two biological replicates. (B) Quantification of (A). Data are presented as mean ± SD (n = 2 biological replicates). (C and D) Representative images (C) and violin plot (D) of γ-H2AX levels in HEK293 cells stably expressing FKBP*-USP11 treated as described in (A). Data are representative of two biological replicates. (E) Venn-diagram representing the number of proteins identified by LC-MS/MS that interact with FKBP*-USP11-FLAG and/or USP11-FLAG, as determined by FLAG pulldown of the indicated FLAG-tagged construct from HEK293T cell lysate.

Additionally, the interactome of FKBP*-USP11 was probed to assess if the FKBP* tag significantly impacts its interaction network. To do this, FKBP*-USP11-FLAG was immunoprecipitated from a cell lysate using anti-FLAG antibody. The precipitated proteins were eluted from the matrix and, after trypsinization, the resultant peptide mixture was analyzed by mass spectrometry. An equivalent experiment was done with USP11-FLAG for comparison. Across three biological replicates, 231 out of 365 proteins co-precipitated with both FKBP*-USP11-FLAG and USP11-FLAG, of which several are known to be involved in DNA DSB repair^38,39^. USP7 and EIF4B, which have previously been reported to associate with USP11^40,41^, were also pulled down (Figure 2E, Data S1).

Taken together, these data argue that FKBP*-USP11 retains some of the known functions of USP11, suggesting that it is a reasonable surrogate for the endogenous protein.

### USP4, USP15 and UCHL1 fusion proteins are degraded by dTAG-13 less efficiently

Next, the degradability of the FKBP*-USP4, -USP15 and -UCHL1 fusions was evaluated. In contrast to USP11, there was only moderate degradation of these constructs after four hours across different concentrations of dTAG-13 (Figures 3A, 3C and 3E). In cells treated with 500 nM dTAG-13, FKBP*-USP4 levels were about 63 % that of vehicle-treated cells (Figure 3A). For FKBP*-USP15 and FKBP*-UCHL1, protein levels of dTAG-13-treated cells were about 65 % and 47 % that of vehicle-treated cells respectively (Figures 3C and 3E). A time course experiment revealed slow degradation of all three constructs (Figures 3B, 3D and 3F). Both FKBP*-USP4 and FKBP*-USP15 levels were greater than 80 % that of vehicle-treated cells after two hours, in stark contrast to the rapid degradation of FKBP*-USP11. Only minimal degradation of FKBP*-UCHL1 was observed two hours after dTAG-13 treatment. More significant degradation was observed at longer times. After 24 hours, FKBP*-USP4, FKBP*-USP15 and FKBP*-UCHL1 levels had dropped to around 9 %, 23 % and 28 %, respectively, of that observed in vehicle-treated cells (Figures 3G – 3L). Overall, these data indicate that the FKBP* fusions to USP4, USP15 and UCHL1 can be degraded by PROTACs but less rapidly and efficiently than FKBP*-USP11.

**Figure 3.**
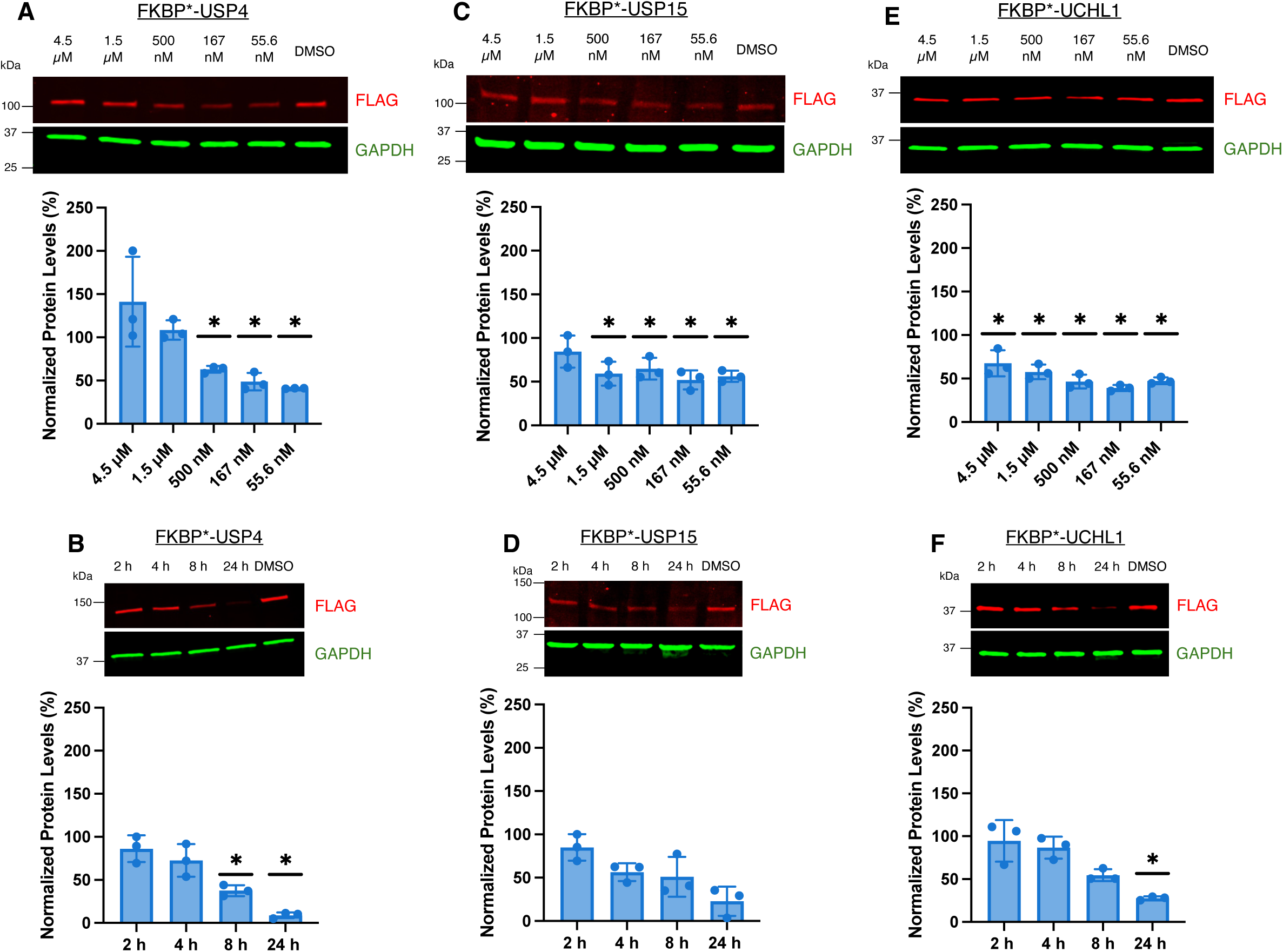
USP4, USP15 and UCHL1 fusion proteins are degraded by dTAG-13 less efficiently. (A, C and E) Representative western blot and quantification of FKBP*-USP4 (A), FKBP*-USP15 (C) or FKBP*-UCHL1 (E) levels after HEK293 cells stably expressing the indicated FKBP* construct were treated for 4 h with a range of dTAG-13 concentrations (55.6 nM – 4.5 µM). Data are presented as mean ± SD (n = 3 biological replicates), unpaired t test with vehicle-treated cells (*q < 0.01). (B, D and F) Representative western blot and quantification of FKBP*-USP4 (B), FKBP*-USP15 (D) or FKBP*-UCHL1 (F) levels after HEK293 cells stably expressing the indicated FKBP* construct were treated with 500 nM dTAG-13 for 2 h, 4 h, 8 h or 24 h. Data are presented as mean ± SD (n = 3 biological replicates), unpaired t test with vehicle-treated cells (*q < 0.01).

### FKBP*-USP4 and -USP15 actively resist degradation through auto-deubiquitylation

To investigate whether the less robust degradation of USP4, USP15 and UCHL1 was due to auto-deubiquitylation, we made mutant constructs in which the catalytic cysteine was replaced with a serine or alanine residue (FKBP*-USP4(C311S), FKBP*-USP15(C298A) and FKBP*-UCHL1(C90A)). These mutant DUBs were then also stably expressed in Flp-In 293 cells.

Treatment of both FKBP*-USP4(C311S) and FKBP*-UCHL1(C90A)-expressing cells with dTAG-13 resulted in greater depletion of the mutant construct compared to wild-type DUB across all the concentrations tested (Figures 4A and 4C). At 500 nM dTAG-13, mutant FKBP*-USP4(C311S) levels were about 8 % of that in vehicle-treated cells. In contrast the level of the wild-type fusion protein was reduced to 63% of that seen in vehicle-treated cells under the same conditions. Similarly, mutant FKBP*-USP15(C298A) levels were about 19% of that in vehicle-treated cells, compared to 65 % for the wild-type construct. We also observed a more rapid rate of degradation of both mutant fusions (Figures 4B and 4D). After two hours, FKBP*-USP4(C311S) levels were approximately 39 % of that observed in vehicle-treated cells, whereas levels of wild-type FKBP*-USP4 were about 86 % of the vehicle control. For FKBP*-USP15(C298A), protein levels were about 27 % of that in vehicle-treated cells compared to about 85 % for wild-type FKBP*-USP15. These data argue that catalytically active FKBP*-USP4 and FKBP*-USP15 resist PROTAC-mediated degradation through auto-deubiquitylation.

**Figure 4.**
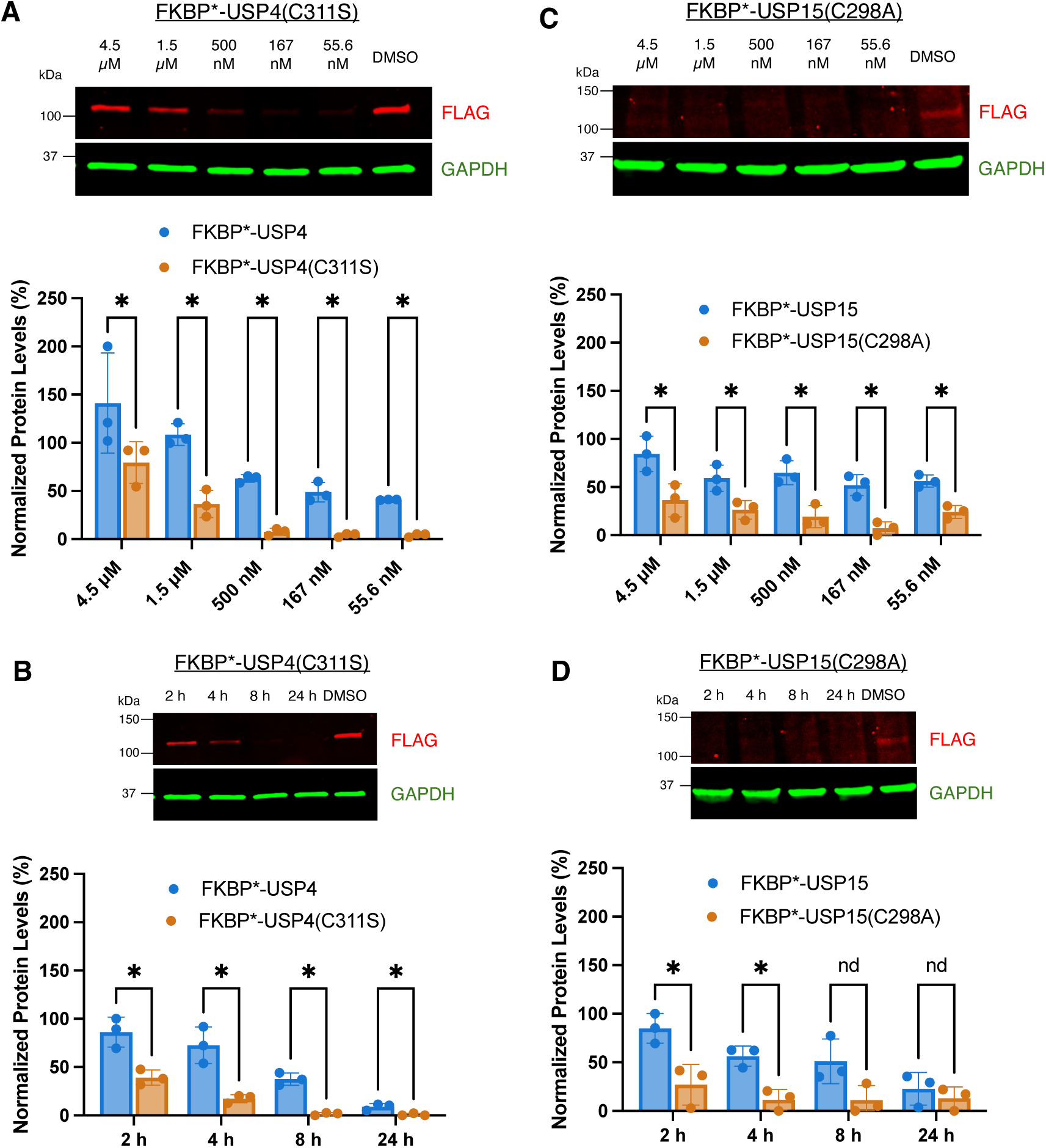
FKBP*-USP4 and -USP15 actively resist degradation through auto-deubiquitylation. (A and C) Representative western blot of FKBP*-USP4(C311S) (A) or FKBP*-USP15(C298A) (C) levels after HEK293 cells stably expressing the indicated FKBP* construct were treated for 4 h with a range of dTAG-13 concentrations (55.6 nM – 4.5 µM). Mutant DUB fusion levels are compared to the wild-type FKBP*-DUB construct (wild-type FKBP*-DUB data are first shown in Figure 3). Data are presented as mean ± SD (n = 3 biological replicates), unpaired t test with Welch correction (*q < 0.05). (B and D) Representative western blot of FKBP*-USP4(C311S) (B) or FKBP*-USP15(C298A) (D) levels after HEK293 cells stably expressing the indicated FKBP* construct were treated with 500 nM dTAG-13 for 2 h, 4 h, 8 h or 24 h. Mutant DUB fusion levels are compared to the wild-type FKBP*-DUB construct (wild-type FKBP*-DUB data are first shown in Figure 3). Data are presented as mean ± SD (n = 3 biological replicates), unpaired t test with Welch correction (*q < 0.05, nd: q ≥ 0.05).

### UCHL1 is degraded slowly because it is a poor proteasome substrate

When FKBP*-UCHL1(C90A)-expressing cells were treated with dTAG-13, we observed that after four hours the levels of the wild-type and mutant fusion proteins were quite similar (Figure 5A). The time course also revealed a comparable rate of degradation between the wild type and catalytically inactive DUB (Figure 5B). Hence, unlike USP4 and USP15, auto-deubiquitylation does not seem to be the main factor contributing to poor degradation of UCHL1.

**Figure 5.**
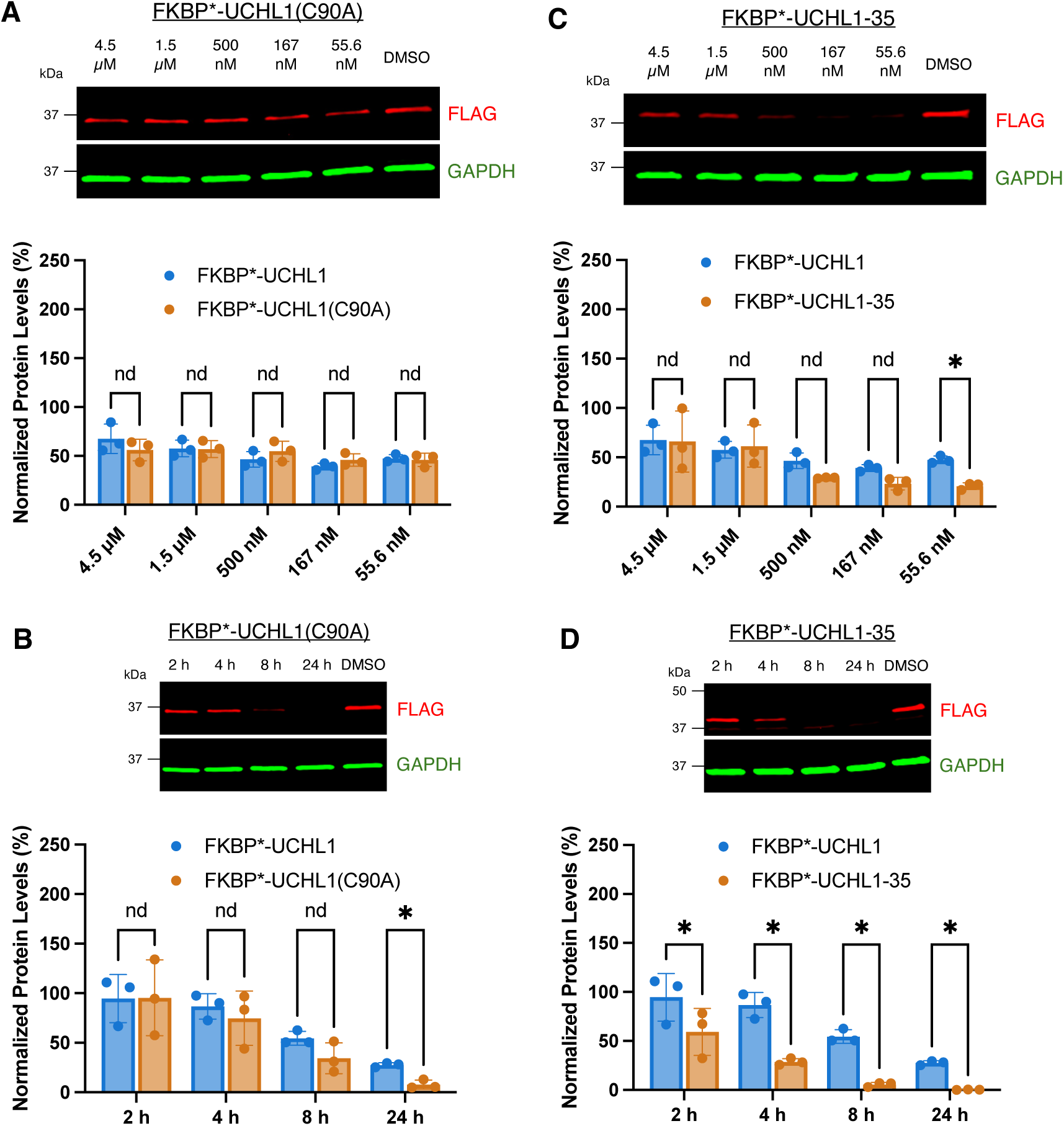
UCHL1 is degraded slowly because it is a poor proteasome substrate. (A and C) Representative western blot of FKBP*-UCHL1(C90A) (A) or FKBP*-UCHL1-35 (C) levels after HEK293 cells stably expressing the indicated FKBP* construct were treated for 4 h with a range of dTAG-13 concentrations (55.6 nM – 4.5 µM). Protein levels are compared to FKBP*-UCHL1 (FKBP*-UCHL1 data are first shown in Figure 3). Data are presented as mean ± SD (n = 3 biological replicates), unpaired t test with Welch correction (*q < 0.05, nd: q ≥ 0.05). (B and D) Representative western blot of FKBP*-UCHL1(C90A) (B) or FKBP*-UCHL1-35 (D) levels after HEK293 cells stably expressing the indicated FKBP* construct were treated with 500 nM dTAG-13 for 2 h, 4 h, 8 h or 24 h. Protein levels are compared to FKBP*-UCHL1 (FKBP*-UCHL1 data are first shown in Figure 3). Data are presented as mean ± SD (n = 3 biological replicates), unpaired t test with Welch correction (*q < 0.05, nd: q ≥ 0.05).

An alternative reason for the poor degradability of FKBP*-UCHL1 could be the lack of an unstructured tail on this protein, a feature important for rapid turnover of proteins by the proteasome.^42,43^ An unstructured region allows facile engagement of the substrate by the proteasomal ATPases that mediate protein unwinding and feed the resultant unstructured chain into the catalytic cavity of the complex. UCHL1 is a structurally characterized, tightly folded protein lacking such an unstructured region^44^.

To test this hypothesis, a 35 amino acid disordered region derived from *S. cerevisiae* Cytochrome b_2_ was added to the C-terminus of FKBP*-UCHL1. This tail has been shown previously to significantly improve the degradability of hard-to-unwind substrates^45^. Indeed, this FKBP*-UCHL1-35 construct was degraded more efficiently than the parent construct. For example, at 55.6 nM dTAG-13, the levels of FKBP*-UCHL1 and FKBP*-UCHL1-35, were 47% and 21%, respectively, of that observed in vehicle-treated cells (Figure 5C). The time course also revealed that the FKBP*-UCHL1-35 construct was degraded more quickly. After just two hours, the level of FKBP*-UCHL1-35 was about 50 % that of vehicle-treated cells, compared to 95 % for unmodified FKBP*-UCHL1 (Figure 5D). These data strongly suggest that the lack of a disordered tail is a significant contributor to sluggish FKBP*-UCHL1 degradation.

To further probe this model, we examined the effect of a p97 inhibitor on the turnover of FKBP*-UCHL1. For protein substrates that lack a disordered tail, the p97 unfoldase complex is responsible for generating the required unstructured region for proteasomal engagement^46,47^. We therefore reasoned that the slower, but measurable, degradation of FKBP*-UCHL1 might require p97 activity, whereas the more rapid degradation of FKBP*-UCHL1-35 may not. Thus, we co-treated FKBP*-UCHL1- and FKBP*-UCHL1-35-expressing cells with dTAG-13 and the p97 inhibitor CB-5083 and assessed the levels of FKBP* fusion protein. As expected, FKBP*-UCHL1 degradation was abolished by the addition of CB-5083, indicating a strong dependence on p97 for degradation (Figures 6A and 6C). Unexpectedly however, the degradation of FKBP*-UCHL1-35 was also abolished when CB-5083 was added, suggesting that the degradation of FKBP*-UCHL1-35 is also dependent on p97 (Figures 6B and 6C). While tangential to the main point of this study, this finding challenges the current paradigm that the main role of p97 is in degrading tightly folded proteins and that it is not essential for turnover of substrates with an intrinsically disordered tail.^46–48^ This is considered further in the discussion section.

**Figure 6.**
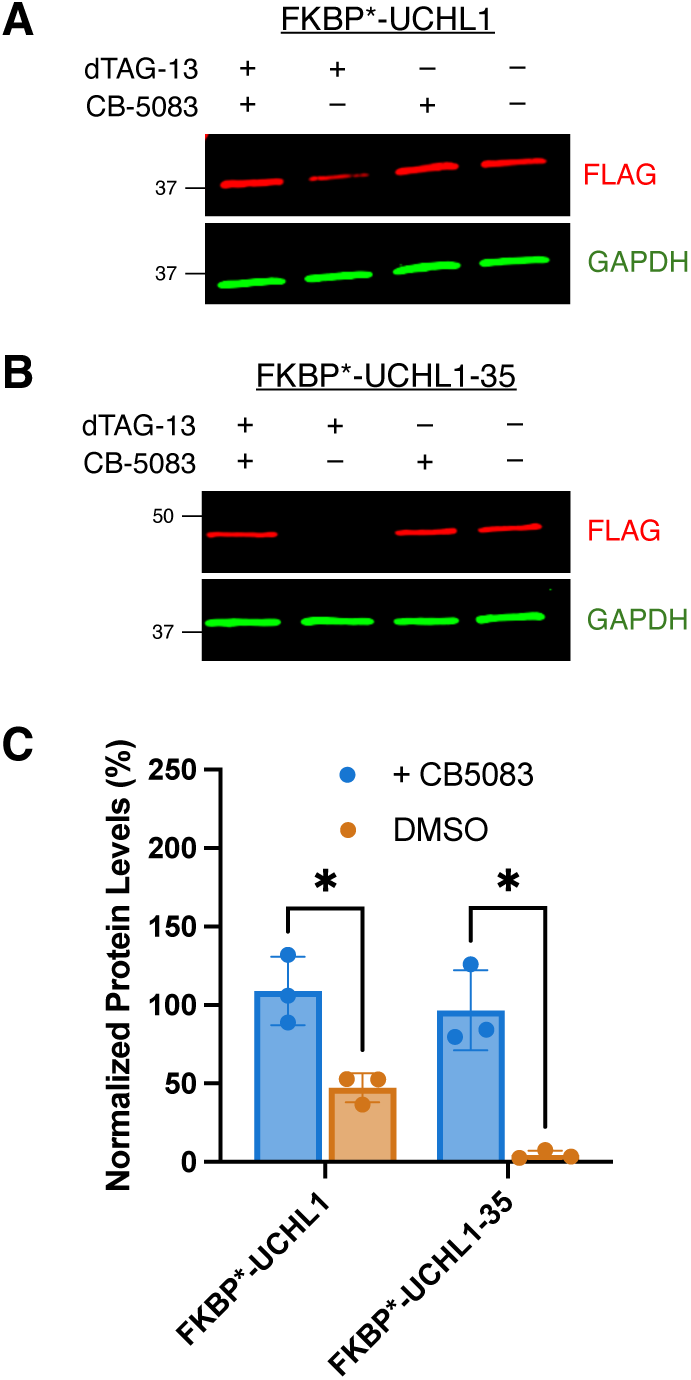
The degradation of both FKBP*-UCHL1 and FKBP*-UCHL1-35 is dependent on p97. (A and B) Western blot of FKBP*-UCHL1 (A) or FKBP*-UCHL1-35 (B) levels after HEK293 cells stably expressing the indicated FKBP* construct were treated with 500 nM dTAG-13 ± 10 µM CB-5083 for 8 h. Blots are representative of three biological replicates. (C) Quantification of (A) and (B). Data are presented as mean ± SD (n = 3 biological replicates), unpaired t test with Welch correction (*q < 0.05).

## Discussion

In this work, we employed dTAG technology^20^ to evaluate the degradability of the DUBs USP11, USP4, UCHL1 and USP15 when engaged by a PROTAC that does not block the active site. Based on our results, we propose three main categories for DUBs with respect to their suitability as targets of PROTAC-style degraders. The first are DUBs that, like USP11, are highly degradable. We found that FKBP*-USP11 levels are reduced rapidly to <10% that observed in vehicle-treated cells (Figure 1). The second comprises DUBs, such as USP4 and USP15, that actively resist degradation through auto-deubiquitylation. These fusion proteins were degraded more slowly and less efficiently than USP11. However, inactivation of their protease activity through mutation of the active site cysteine rendered them more susceptible to degradation (Figures 3 and 4). The third category includes UCHL1, which is degraded inefficiently (Figure 3) but not because of its catalytic activity. In this case, the protein is simply a poor proteasome substrate due to the absence of an unstructured tail that can be easily engaged by the AAA class ATPases in the proteasome that are responsible for substrate unfolding. This view is supported by the observation that addition of a 35 amino acid unstructured tail to FKBP*-UCHL1 strongly stimulated its degradation upon exposure of cells to dTAG-13 (Figure 5). Interestingly, for UCHL1 and USP4, the resistance to PROTAC-mediated degradation is consistent with the reported turnover rate of these DUBs in mammalian cells. UCHL1 has been shown to have a half-life of greater than 24 h^49^, while wild-type USP4 was found to be more stable than its catalytically inactive counterpart^50^.

A side note that is peripheral to this investigation, but of interest with respect to the fundamental mechanism of protein turnover in cells, is our observation that even FKBP*-UCHL1-35 is highly dependent on p97 for its degradation (Figure 6). As mentioned above, this runs counter to current dogma in the field that suggests p97 is dispensable for the turnover of proteins with an unstructured tail. This could be explained by UCHL1’s rather unusual structure. Its backbone forms a 5-2 knot^51^, such that pulling on either terminus will still lead to a knotted structure. Thus, even if the AAA-ATPases of the proteasome engage FKBP*-UCHL1-35, the unwinding process is likely to stall and require p97 to unknot the protein.

Returning to the main theme of this study, we believe that this workflow is useful to de-risk a PROTAC development against a particular DUB of interest since the development of high-quality ligands to these targets is a significant undertaking. In this vein, USP11-class DUBs, which are readily degraded, are especially interesting as these are presumably the preferred targets for non-inhibitory PROTACs. However, as is the case in any chemical genetics experiment, care needs to be taken in applying conclusions reached through analysis of a fusion construct to the native protein. In this case, a particular concern is that the fusion of the FKBP mutant to USP11 compromises the catalytic activity of the protein and that its sensitivity to PROTAC-mediated degradation is an artifact.

This seems unlikely for several reasons. First, we demonstrated that FKBP*-USP11 is able to substitute for the endogenous protein in the repair of DNA double-strand breaks, as monitored by an accumulation of γ-H2AX foci that are a marker of unrepaired breaks (Fig. 2). The DUB activity of USP11 is essential for this process, so this observation argues that FKBP*-USP11 is an active DUB. Second, we showed that FKBP*-USP11 associates with USP7 and EIF4B, known interaction partners of USP11, as well as several components of the DSB repair complex such as PRKDC and XRCC5. We had considered also evaluating immunoprecipitated FKBP*-USP11 for DUB activity using a fluorogenic assay but this is complicated by the presence of other DUBs, including USP7 in the immunoprecipitate. Finally, USP11 is a paralog of USP4 and USP15, sharing 45 % and 46 % identity with these proteins, respectively. We found that both USP4 and USP15 resist turnover through auto-deubiquitylation, so it seems unlikely that the identical fusion to USP11 would compromise its catalytic activity. Taken together, these data strongly suggest that FKBP*-USP11 is an active DUB. However, we acknowledge that it is impossible to say unequivocally that the sensitivity of FKBP*-USP11 to PROTAC-mediated turnover will indeed reflect that of native USP11 once appropriate ligands for the construction of non-inhibitory PROTACs become available.

The significant difference observed in the degradability of the FKBP*-USP11, -USP4 and - USP15 is interesting from the point of view of developing selective degraders of USP11. These proteins are very closely related paralogs and it may be difficult to identify compounds that distinguish cleanly between them even using ligands that engage the protein outside the active site. However, the data reported here suggest the possibility that even a PROTAC that engages all three paralogs might display significant selectivity for USP11 degradation if it is the least efficient of three in deubiquitylating itself. There are several examples of large divergences between proteins that are engaged by a PROTAC and those that are degraded^21,22^. But these are attributed to differences in the efficiency of poly-Ubiquitylation of the substrate by the E3 ligase. This issue of more or less efficient auto-deubiquitylation would represent an alternative mechanism for achieving selectivity.

## Limitations of the Study

A limitation of our approach to evaluate the vulnerability of an uninhibited Dub to PROTAC-mediated poly-Ubiquitylation and subsequent turnover is that artificial fusion proteins are employed. This is unavoidable since non-inhibitory ligands for almost all Dubs do not exist. But it is certainly possible that a PROTAC eventually developed against a native Dub might target different lysine residues for modification than is the case using the dTAG system. This, in turn could affect the efficiency of auto-deubiquitylation and thus Dub turnover. Therefore, this platform should be considered as a tool for de-risking a Dub degrader project, not as a strict predictor of the eventual success or failure in the development of a *bona fide* PROTAC.

## Resource Availability

### Lead contact

Requests for further information should be directed to the lead contact, Thomas Kodadek

### Materials availability

All plasmids and stable cell lines generated in this study are available from the lead contact upon request.

### Data and code availability

Mass spectrometry proteomics data have been deposited to the ProteomeXchange Consortium (http://proteomecentral.proteomexchange.org) via the iProX partner repository as iProX: PXD071411 and are publicly available as of the date of publication. This paper does not report original code. Any additional information required to reanalyze the data reported in this paper is available from the lead contact upon request.

## Acknowledgements

This work was supported by grants from the U.S. National Cancer Institute (RO1 CA290247) and the Office of the Director, NIH (S10OD036363). We would like to thank the Mass Spectrometry and Proteomics Core Facility at the Herbert Wertheim UF Scripps Institute for Biomedical Innovation & Technology for assistance with the FKBP*-USP11 interactome experiment.

## Author contributions

Both T.K. and J.T. conceived this study. J.T. performed most of the experiments, with help from J.M.W. for the imaging experiment and the Proteomics Core for the FKBP*-USP11 interactome experiment. Both T.K. and J.T. wrote and edited the manuscript. The entire study was supervised by T.K. and J.M.B.

## Methods Section

### Mammalian Cell lines

Flp-In 293 cells were purchased from ThermoFisher while HEK293T cells were obtained from American Type Culture Collection (ATCC). Both cell lines are of female origin. Flp-In 293 cells and stable Flp-In 293 cell lines expressing FKBP* constructs were grown in in DMEM (4.5 g/L D-Glucose) supplemented with 10 % FBS, 2 mM GlutaMAX and 100 μg/mL Zeocin (Flp-In 293) or 100 μg/mL Hygromycin B (stable Flp-In 293 cell lines). HEK293T cells wesre grown in DMEM (4.5 g/L D-Glucose) supplemented with 10 % fetal bovine serum (FBS), 4 mM L-alanyl-L-glutamine (GlutaMAX) and 110 mg/L sodium pyruvate. Cells were maintained at 37 °C with 5 % CO_2_.

### Bacteria Culture

Stellar competent cells (*E. coli* HST08 strain) were purchased from Takara Bio. To prepare cultures for mini- and maxi-preps, cells were grown in LB + 100 μg/ml Ampicillin and shaken at 37 °C for 16 h.

### Plasmid construction

The pcDNA5-FRT plasmid backbone was PCR amplified with indicated primers (IDT, see table S1) from the pcDNA5-FRT-HaloTag7 plasmid^52^. 1 ng of the template plasmid, forward and reverse primers (final concentration of 300 nM) and 25 μl of PrimeSTAR Max DNA Polymerase (Takara) ws added to Milli-Q water to a final volume of 50 ul. The mixture was then run on a Bio-Rad T100 thermal cycler (30 cycles of 98 °C for 10 s, 55 °C for 10 s, 68 °C for 162 s). The PCR reaction mixture was run on a 1 % agarose gel and the linearized plasmid excised from the gel and extracted with a NucleoSpin Gel and PCR Clean-up kit (Machery-Nagel). gBlocks Gene Fragments encoding FKBP* constructs (including overhangs) were purchased from IDT and inserted into the pcDNA5-FRT plasmid backbone by In-Fusion cloning (Takara). The linearized pcDNA5-FRT vector and purchased Gene Fragments (amount used was determined by the online Takara In-Fusion molar ratio calculator) was added to 2 μl of 5X In-Fusion Snap Assembly Master Mix (Takara) and Milli-Q water to a final volume of 10 μl. The infusion reaction was incubated at 50 °C for 15 minutes on a Bio-Rad T100 thermal cycler. 2.5 μl of the Infusion reaction was then added to 50 ul of Stellar Competent cells (Takara) and incubated on ice for 30 minutes. The cells were then heat shocked at 42 °C for 45 s, followed by a two minute recovery on ice. 450 μl of SOC media was added, and the cells were shaken at 37 °C for 1 h. 50 μl of cells were then plated on LB Agar (Fisher Scientific) + 100 μg/ml Ampicillin (USBiological Life Sciences) and incubated at 37 °C for 16 h. After 16 h, single colonies were picked and grown for 16 h at 37 °C (with shaking) in 5 ml of LB (Fisher Scientific) + 100 μg/ml Ampicillin. Plasmid DNA was extracted using the QIAPrep Spin Miniprep Kit (Qiagen) and sent for full plasmid sequencing (Azenta Life Sciences). For the correct plasmid constructs, bacteria glycerol stocks were inoculated into 200 ml of LB + 100 μg/ml Ampicillin and shaken at 37 °C for 16 h. Purified plasmid DNA was obtained using a HiSpeed Plasmid Maxi Kit (Qiagen). For the mutant FKBP* constructs, the template plasmid was PCR amplified with the indicated mutagenesis primers (IDT, see table S1). 1 ng of the template plasmid, forward and reverse primers (final concentration of 300 nM) and 12.5 μl of PrimeSTAR Max DNA Polymerase (Takara) was added to Milli-Q water to a final volume of 25 ul. The mixture was then run on a Bio-Rad T100 thermal cycler (35 cycles of 98 °C for 10 s, 55 °C for 10 s, 72 °C for 45 s). Linearized DNA was obtained by gel extraction as described earlier, and then re-circularized by In-Fusion cloning (Takara). 100 ng of the linearized DNA was added to 2 μl of 5X In-Fusion Snap Assembly Master Mix (Takara) and Milli-Q water to a final volume of 10 μl, followed by incubation at 50 °C for 15 minutes on a Bio-Rad T100 thermal cycler. Transformation into competent cells, plating, minipreps and maxipreps were then carried out in the same way as the wild type constructs.

### Generation of Flp-In 293 cells stably expressing FKBP* constructs

Flp-In 293 cells were seeded in a 10 cm dish to be about 70 % confluent the next day and incubated at 37 °C overnight. The next day, 21.6 μg of pOG44 plasmid (ThermoFisher) and 2.4 μg of FKBP* construct plasmid was added to Opti-MEM (ThermoFisher) to a total volume of 1.5 ml. 60 μl of Lipofectamine 2000 (ThermoFisher) was then added to 1.44 ml of Opti-MEM and incubated at 25 °C for five minutes. The plasmid mixture was then added to the Lipofectamine mixture dropwise, mixed gently and incubated at 25 °C for 20 minutes. After 20 minutes, media in the 10 cm dish was aspirated and replaced with fresh media. The plasmid-Lipofectamine mixture was added dropwise to the cells, and the cells then incubated at 37 °C for 24 h. The media was refreshed after 24 hours. After an additional 24 hours, cells were split into four 10 cm dishes in media supplemented with 100 μg/mL Hygromycin B (ThermoFisher). After seven days, Hygromycin B-resistant cells were pooled and expanded into four T175 flasks. When the flasks were confluent, the cells were collected and frozen in 10 % DMSO in Flp-In 293 cell media (about 1.5 X 10^7^ cells / cryotube). Cells were slow frozen in a Mr. Frosty freezing container at – 80 °C, and then transferred to – 150 °C for long term storage. Expression of the FKBP* constructs in was validated by Western Blotting (anti-FLAG).

### Degradation assay

Flp-In 293 cells stably expressing FKBP* constructs were seeded in 6-well plates with complete media lacking antibiotics and incubated at 37 °C overnight. The next day, cells were treated with varying concentrations of dTAG-13 (final concentration 4.5 μM – 55.6 nM) and incubated at 37 °C for four hours before collection. For the time course, cells were treated with 500 nM dTAG-13 (final concentration) and incubated at 37 °C for two, four, eight or 24 hours before collection. The cells were pelleted at 500 xg for five minutes and the supernatant was removed before addition of the lysis cocktail.

### Evaluating dependence on the proteasome, ubiquitin or p97 for degradation

Flp-In 293 cells stably expressing FKBP* constructs were seeded in 6-well plates with complete media lacking antibiotics and incubated at 37 °C overnight. The next day, cells were co-treated with a final concentration of 500 nM dTAG-13 and 10 μM Bortezomib (proteasome inhibitor), TAK-243 (ubiquitin activating enzyme UBA1 inhibitor) or CB-5083 (p97 inhibitor) and incubated at 37 °C for four (Bortezomib and TAK-243) or eight hours (CB-5083) before collection. Cells were pelleted at 500 xg for five minutes and the supernatant was removed before addition of the lysis cocktail.

### Western blotting

Cells were lysed in M-PER Mammalian Protein Extraction Reagent + 1X Halt Protease Inhibitor Cocktail (ThermoFisher) (100 μl / sample) for 12 minutes and clarified by centrifugation at 14,000 xg for 15 minutes. Protein concentration was determined by Bradford assay and normalized gel samples were prepared with 4X Laemmli sample buffer (Bio-Rad) supplemented with 2-mercaptoethanol, DPBS and cell lysate (total volume of 66.7 μl). Gel samples were loaded on 4 – 20 % TGX gels (Bio-Rad) and run at 180 V for 35 – 40 minutes. Gels were transferred to a nitrocellulose membrane (Bio-Rad) using the TransBlot Turbo semi-dry transfer system (mixed MW setting). Membranes were blocked with 5 % Blot-QuickBlocker (G-Biosciences) in PBST for one hour and then incubated with primary antibodies (see key resources table) in the same blocking buffer at 4 °C overnight. The membranes were then washed with PBST (3 X 15 min) and incubated with secondary antibodies (see key resources table) in Intercept Blocking Buffer (Li-Cor) at 25 °C for one hour. Membranes were washed again with PBST (3 X 15 min) and imaged on a Li-Cor Odyssey M scanner. All incubation and washing steps with the membrane were done on an orbital shaker.

### USP11 depletion for γ-H2AX assay

Flp-In 293 cells stably expressing FKBP*-USP11 were seeded in 6-well plates (to be about 70 % confluent the next day) with complete media lacking antibiotics and incubated at 37 °C overnight. The next day, 5 μM siRNA solutions were prepared from the 20 μM stock solution. 20 μl of 5 μM siRNA solution was added to 180 μl of Opti-MEM (ThermoFisher) and mixed gently. 5 μl of DharmaFECT1 (Horizon Discovery) was then added to 195 μl of Opti-MEM, mixed gently and incubated at 25 °C for five minutes. The siRNA mixture was then added to the DharmaFECT1 mixture dropwise, mixed gently and incubated at 25 °C for 20 minutes. After 20 minutes, media in the 6-well plate was aspirated and replaced with fresh media. The siRNA-DharmaFECT1 mixture was added dropwise to the cells, and the cells then incubated at 37 °C for 24 h. Media was refreshed after 24 h. After an additional 24 hours of incubation at 37 °C, 15 mm round coverslips were placed into a half of the wells in a 12-well plate and pre-treated with 70 % ethanol for two minutes at 25 °C and then aspirated and air-dried under the TC hood. 1 ml of Poly-L-Lysine solution (Sigma-Aldrich) was then added to each well, left for five minutes 37 °C and then aspirated and air-dried. The cells were then split into the 12-well plate, with two wells for each experimental condition (one for imaging, one for Western Blot analysis), and treated with a final concentration of 500 nM dTAG-13 or vehicle control. After 24 hours of incubation at 37 °C, the cells were either collected for western blot analysis (pelleted at 500 xg for five minutes) or used for immunofluorescent staining.

### Immunofluorescence imaging

Cells in a 12-well plate were washed with PBS (X1) and fixed with 4 % formaldehyde in PBS for 10 minutes at 25 °C. The cells were then washed with PBS (X1) and permeabilized with cold 70 % ethanol for two hours at 4 °C. Cells were washed with PBS (X3) and then incubated with γ-H2AX antibody (Cell Signaling Technology) in 1 % BSA (PBS) at 4 °C overnight. The cells were then washed with PBS (3 X 15 min) and incubated with α-rabbit secondary antibody (Abcam) in in 1 % BSA (PBS) at 4 °C for two hours. Cells were then washed again with PBS (3 X 15 min). Cover slips were mounted onto glass slides with VECTASHIELD PLUS mounting medium with DAPI (Vector Laboratories) and dried. Cells were imaged on a Nikon Eclipse Ti2 with a CFI60 Plan Apochromat Lambda D 100x Oil Immersion Objective Lens, N.A. 1.45, W.D. 0.13mm, F.O.V. 25mm, DIC, Spring Loaded.

### FKBP*-USP11-FLAG / USP11-FLAG pulldown

HEK293T cells were seeded in a 6-well plate with complete media lacking antibiotics and incubated at 37 °C overnight. The next day, 0.85 μg of FKBP*-USP11-FLAG or USP11-FLAG plasmid was added to Opti-MEM (ThermoFisher) to a total volume of 250 μl. 8 μl of Lipofectamine 2000 (ThermoFisher) was then added to 242 μl of Opti-MEM and incubated at 25 °C for five minutes. The plasmid mixture was then added to the Lipofectamine mixture dropwise, mixed gently and incubated at 25 °C for 20 minutes. After 20 minutes, media in the 6-well plate was aspirated and replaced with fresh media. The plasmid-Lipofectamine mixture was added dropwise to the cells, and the cells then incubated at 37 °C for 24 h. After 24 h, the cells were collected and pelleted at 500 xg for five minutes. Cells were then lysed and clarified (see method details for Western Blotting). Protein concentration was determined by Bradford assay and 4 mg/ml samples (200 μl each) were prepared with TBS. 60 ul of anti-FLAG M2 magnetic beads (Millipore) was equilibrated into TBS (3X washes with 200 μl TBS). 150 μl of 4 mg/ml lysate-TBS mixture was added to the beads and tumbled at 4 °C for two hours. The beads were washed with TBS (3 X 400 ul) and proteins were eluted with 150 μl 2X Laemmli sample buffer (Bio-Rad) + 150 μl TBS. Eluted proteins were loaded on 4 – 20 % TGX gels (Bio-Rad) and run at 180 V for about five minutes. The band was excised from the gel and used for LC-MS/MS analysis.

### LC-MS/MS

Gel fractionated proteins were in-gel digested with trypsin (Pierce Biotechnology) for 1 hour at 50 °C using ProteaseMax^TM^ Surfactant trypsin enhancer following reduction and alkylation with dithiothreitol and iodoacetamide, respectively, according to the manufacturer’s instructions (Promega). LC-MS/MS analysis of extracted peptides was subsequently carried out using an Orbitrap Fusion Tribrid mass spectrometer, following 2 mg capacity ZipTip (Millipore) C18 sample clean-up according to the manufacturer’s instructions. Peptides were eluted from an EASY PepMapTM RSLC C18 column (Thermo) into the mass spectrometer using a gradient of 5 – 25 % solvent B (80/20 acetonitrile/water, 0.1% formic acid) in 90 minutes, followed by 25 – 44 % solvent B in 30 minutes, 44 – 80 % solvent B in 0.10 minute, a 10 minute-hold of 80 % solvent B, a return to 5 % solvent B in 3 minutes, and finally with another 3-minute hold of 5 % solvent B. The gradient was then extended for the purpose of cleaning the column by increasing solvent B to 98 % in 3 minutes, a 98 % solvent B hold for 10 minutes, a return to 5 % solvent B in 3 minutes, a 5 % solvent B fold for 3 minutes, an increase of solvent B to 98 % in 3 minutes, a 98 % solvent B hold for 10 minutes, a return to 5 % solvent B in 3 minutes and a 5 % solvent B hold for 3 minutes and finally, another increase to 98 % solvent B in 3 minutes and a hold of 98 % solvent B for 10 minutes. All flow rates were 250 nL/min delivered using a Vanquish Neo UHPLC (Thermo Fisher Scientific, San Jose, CA). Solvent A consisted of 0.1% formic acid. Ions were created with an EASY Spray source (Thermo Scientific, San Jose, CA) held at 45oC using a voltage of 2.2kV. Data dependent scanning was performed by the Xcalibur v 4.7.69.37 software using a survey scan at 120, 000 resolution in the Orbitrap analyzer scanning mass/charge (m/z) 350-2000 followed by higher-energy collisional dissociation (HCD) tandem mass spectrometry (MS/MS) at a normalized collision energy of 30% of the most intense ions at maximum speed, at an automatic gain control of 1.0E4. Precursor ions were selected by the monoisotopic precursor selection (MIPS) setting to peptide and MS/MS was performed on charged species of 2-8 charges at a resolution of 30,000. Dynamic exclusion was set to exclude ions once within a 25 second window. All scan events occurred within a 2-second specified cycle time.

### Protein/peptide identification

Tandem mass spectra were searched against the human proteome protein sequences from Uniprot (UP000005640) downloaded on July 09, 2023, and common contaminant proteins available with Proteome Discoverer v 2.5.0.400. At the time of the search the UP0000005640 contained 20523 sequences, and the contaminant protein database contained an additional 298 sequences. All MS/MS spectra were searched using Thermo Proteome Discoverer 2.5.0.400 (Thermo) considering fully tryptic peptides with up to 2 missed cleavage sites. Variable modifications considered during the search included methionine oxidation (15.995 Da), and asparagine and qlutamine deamidation (0.984 Da). Cysteine carbamidomethylation (57.021 Da) was considered as a static modification. Proteins were identified at 99% confidence with XCorr score cut-offs^53^ as determined by a reversed database search. The Minora Feature Detector node which detects chromatographic peaks and features as also used in Proteome Discoverer at default settings. The protein and peptide identification results were also visualized with Scaffold v 5.0.0 (Proteome Software Inc), a program that relies on various search engine results (i.e.: Sequest, X!Tandem, MASCOT) and which uses Bayesian statistics to reliably identify more spectra^54^. Proteins were accepted that passed a minimum of two peptides identified at 1% peptide and protein FDR, within Scaffold. The mass spectrometry analysis was performed at The Herbert Wertheim UF Scripps Institute for Biomedical Innovation & Technology, Mass Spectrometry and Proteomics Core Facility (RRID:SCR_023576).

### Quantification and Statistical Analysis

Western blots were quantified using the Emperia Studio software (Li-Cor). Statical analysis was performed with Graphpad Prism 10; details can be found in the figure legends. For the microscopy experiment, two fields of view were used for quantification. γ-H2AX signal for each cell was quantified using Fiji software. For analysis of tandem mass spec data, details can be found in the method details (under protein/peptide identification).

---

**Figure S1.**
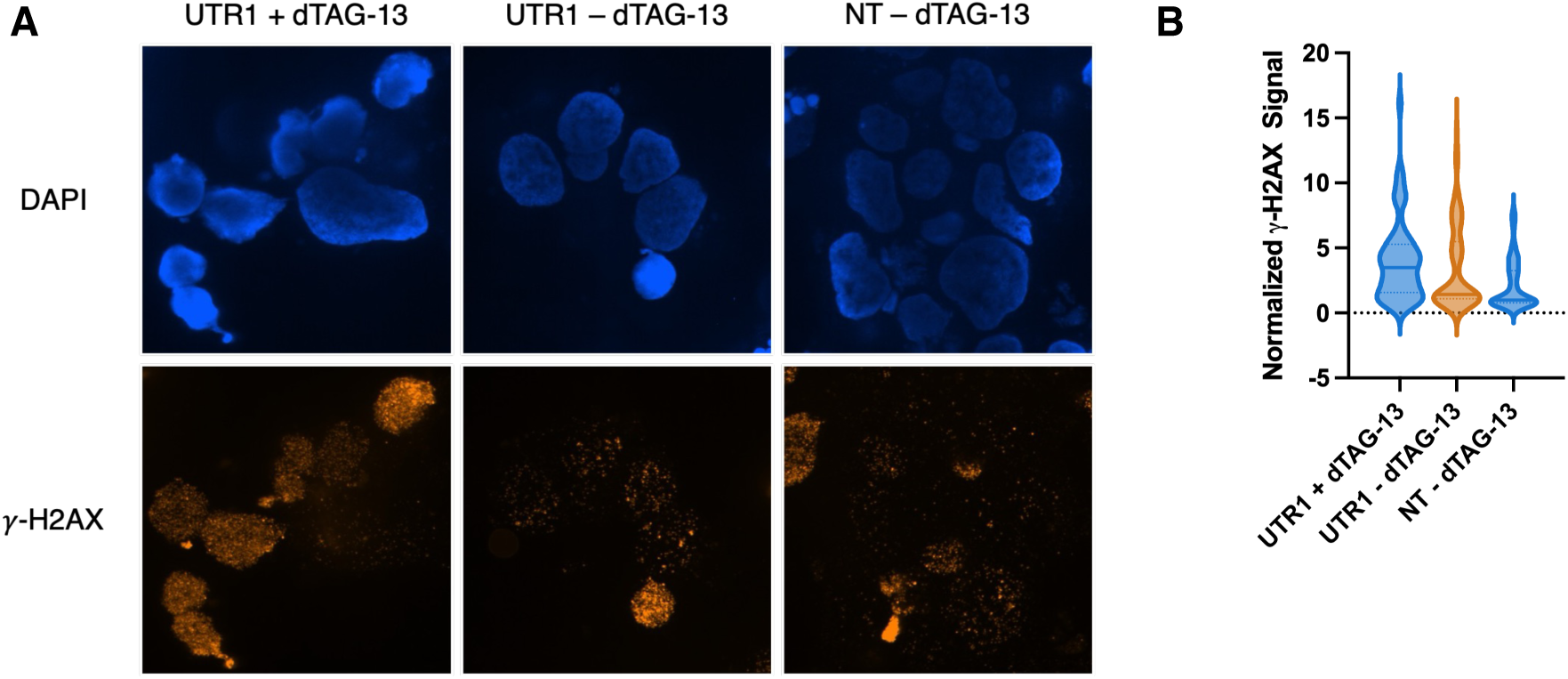
Microscopy data for the second biological replicate of experiment shown in Figure 2C and D. (A and B) Representative images (A) and violin plot (B) of γ-H2AX levels in HEK293 cells stably expressing FKBP*-USP11 treated as described in Figure 2A.

## References

1. Rape, M. (2018). Ubiquitylation at the crossroads of development and disease. Nat. Rev. Mol. Cell. Biol .19, 59–70.

2. Lange, S.M., Armstrong, L.A., and Kulathu, Y. (2022). Deubiquitinases: From mechanisms to their inhibition by small molecules. Mol Cell 82, 15–29.

3. Li, M., Brooks, C.L., Kon, N., and Gu, W. (2004). A Dynamic Role of HAUSP in the p53-Mdm2 Pathway. Mol. Cell 13, 879–886.

4. Cummins, J.M., Rago, C., Kohli, M., Kinzler, K.W., Lengauer, C., and Vogelstein, B. (2004). Tumour suppression: disruption of HAUSP gene stabilizes p53. Nature 428, 1–2.

5. Pornour, M., Jeon, H.Y., Ryu, H., Khadka, S., Xu, R., Chen, H., Hussain, A., Lam, H.M., Zhuang, Z., Oo, H.Z., et al. (2024). USP11 promotes prostate cancer progression by up-regulating AR and c-Myc activity. Proc. Natl. Acad. Sci. U S A 121, e2403331121.

6. Zhang, N., Wang, Q., Lu, Y., Wang, F., and He, Z. (2024). The deubiquitinating enzyme USP11 regulates breast cancer progression by stabilizing PGAM5. Breast Cancer Res 26, 135.

7. Bingol, B., Tea, J.S., Phu, L., Reichelt, M., Bakalarski, C.E., Song, Q., Foreman, O., Kirkpatrick, D.S., and Sheng, M. (2014). The mitochondrial deubiquitinase USP30 opposes parkin-mediated mitophagy. Nature 510, 370–375.

8. Lee, B.H., Lee, M.J., Park, S., Oh, D.C., Elsasser, S., Chen, P.C., Gartner, C., Dimova, N., Hanna, J., Gygi, S.P., et al. (2010). Enhancement of proteasome activity by a small-molecule inhibitor of USP14. Nature 467, 179–184.

9. Turnbull, A.P., Ioannidis, S., Krajewski, W.W., Pinto-Fernandez, A., Heride, C., Martin, A.C.L., Tonkin, L.M., Townsend, E.C., Buker, S.M., Lancia, D.R., et al. (2017). Molecular basis of USP7 inhibition by selective small-molecule inhibitors. Nature 550, 481–486.

10. Kategaya, L., Di Lello, P., Rouge, L., Pastor, R., Clark, K.R., Drummond, J., Kleinheinz, T., Lin, E., Upton, J.P., Prakash, S., et al. (2017). USP7 small-molecule inhibitors interfere with ubiquitin binding. Nature 550, 534–538.

11. Schauer, N.J., Liu, X., Magin, R.S., Doherty, L.M., Chan, W.C., Ficarro, S.B., Hu, W., Roberts, R.M., Iacob, R.E., Stolte, B., et al. (2020). Selective USP7 inhibition elicits cancer cell killing through a p53-dependent mechanism. Sci. Rep. 10, 5324.

12. Clancy, A., Heride, C., Pinto-Fernandez, A., Elcocks, H., Kallinos, A., Kayser-Bricker, K.J., Wang, W., Smith, V., Davis, S., Fessler, S., et al. (2021). The deubiquitylase USP9X controls ribosomal stalling. J. Cell Biol. 220.

13. Fang, T.Z., Sun, Y., Pearce, A.C., Eleuteri, S., Kemp, M., Luckhurst, C.A., Williams, R., Mills, R., Almond, S., Burzynski, L., et al. (2023). Knockout or inhibition of USP30 protects dopaminergic neurons in a Parkinson’s disease mouse model. Nat. Commun. 14, 7295.

14. Chan, W.C., Liu, X., Magin, R.S., Girardi, N.M., Ficarro, S.B., Hu, W., Tarazona Guzman, M.I., Starnbach, C.A., Felix, A., Adelmant, G., et al. (2023). Accelerating inhibitor discovery for deubiquitinating enzymes. Nat. Commun. 14, 686.

15. Meray, R.K., and Lansbury, P.T., Jr. (2007). Reversible monoubiquitination regulates the Parkinson disease-associated ubiquitin hydrolase UCH-L1. J. Biol. Chem. 282, 10567–10575.

16. Denuc, A., Bosch-Comas, A., Gonzalez-Duarte, R., and Marfany, G. (2009). The UBA-UIM domains of the USP25 regulate the enzyme ubiquitination state and modulate substrate recognition. PLoS One 4, e5571.

17. Wijnhoven, P., Konietzny, R., Blackford, A.N., Travers, J., Kessler, B.M., Nishi, R., and Jackson, S.P. (2015). USP4 Auto-Deubiquitylation Promotes Homologous Recombination. Mol. Cell 60, 362–373.

18. Shen, C., Ye, Y., Robertson, S.E., Lau, A.W., Mak, D.O., and Chou, M.M. (2005). Calcium/calmodulin regulates ubiquitination of the ubiquitin-specific protease TRE17/USP6. J. Biol. Chem. 280, 35967–35973.

19. Peng, Y., Liao, Q., Tan, W., Peng, C., Hu, Z., Chen, Y., Li, Z., Li, J., Zhen, B., Zhu, W., et al. (2019). The deubiquitylating enzyme USP15 regulates homologous recombination repair and cancer cell response to PARP inhibitors. Nat. Commun. 10, 1224.

20. Nabet, B., Roberts, J.M., Buckley, D.L., Paulk, J., Dastjerdi, S., Yang, A., Leggett, A.L., Erb, M.A., Lawlor, M.A., Souza, A., et al. (2018). The dTAG system for immediate and target-specific protein degradation. Nat. Chem. Biol. 14, 431–441.

21. Huang, H.T., Dobrovolsky, D., Paulk, J., Yang, G., Weisberg, E.L., Doctor, Z.M., Buckley, D.L., Cho, J.H., Ko, E., Jang, J., et al. (2018). A Chemoproteomic Approach to Query the Degradable Kinome Using a Multi-kinase Degrader. Cell Chem. Biol. 25, 88–99 e86.

22. Bondeson, D.P., Smith, B.E., Burslem, G.M., Buhimschi, A.D., Hines, J., Jaime-Figueroa, S., Wang, J., Hamman, B.D., Ishchenko, A., and Crews, C.M. (2018). Lessons in PROTAC Design from Selective Degradation with a Promiscuous Warhead. Cell Chem. Biol. 25, 78–87 e75.

23. Donovan, K.A., Ferguson, F.M., Bushman, J.W., Eleuteri, N.A., Bhunia, D., Ryu, S., Tan, L., Shi, K., Yue, H., Liu, X., et al. (2020). Mapping the Degradable Kinome Provides a Resource for Expedited Degrader Development. Cell 183, 1714–1731.

24. Bensimon, A., Pizzagalli, M.D., Kartnig, F., Dvorak, V., Essletzbichler, P., Winter, G.E., and Superti-Furga, G. (2020). Targeted Degradation of SLC Transporters Reveals Amenability of Multi-Pass Transmembrane Proteins to Ligand-Induced Proteolysis. Cell Chem. Biol. 27, 728–739 e729.

25. Tian, X., Isamiddinova, N.S., Peroutka, R.J., Goldenberg, S.J., Mattern, M.R., Nicholson, B., and Leach, C. (2011). Characterization of selective ubiquitin and ubiquitin-like protease inhibitors using a fluorescence-based multiplex assay format. Assay Drug Dev. Technol. 9, 165–173.

26. Conole, D., Cao, F., Am Ende, C.W., Xue, L., Kantesaria, S., Kang, D., Jin, J., Owen, D., Lohr, L., Schenone, M., et al. (2023). Discovery of a Potent Deubiquitinase (DUB) Small-Molecule Activity-Based Probe Enables Broad Spectrum DUB Activity Profiling in Living Cells. Angew. Chem. Int. Ed. Engl. 62, e202311190.

27. Zhang, H., Han, Y., Xiao, W., Gao, Y., Sui, Z., Ren, P., Meng, F., Tang, P., and Yu, Z. (2023). USP4 promotes the proliferation, migration, and invasion of esophageal squamous cell carcinoma by targeting TAK1. Cell Death Dis. 14, 730.

28. Zhang, X., Berger, F.G., Yang, J., and Lu, X. (2011). USP4 inhibits p53 through deubiquitinating and stabilizing ARF-BP1. EMBO J. 30, 2177–2189.

29. Chen, X.S., Wang, K.S., Guo, W., Li, L.Y., Yu, P., Sun, X.Y., Wang, H.Y., Guan, Y.D., Tao, Y.G., Ding, B.N., et al. (2020). UCH-L1-mediated Down-regulation of Estrogen Receptor alpha Contributes to Insensitivity to Endocrine Therapy for Breast Cancer. Theranostics 10, 1833–1848.

30. Yao, J., Reyimu, A., Sun, A., Duoji, Z., Zhou, W., Liang, S., Hu, S., Wang, X., Dai, J., and Xu, X. (2022). UCHL1 acts as a potential oncogene and affects sensitivity of common anti-tumor drugs in lung adenocarcinoma. World J. Surg. Oncol. 20, 153.

31. Chen, L.L., Smith, M.D., Lv, L., Nakagawa, T., Li, Z., Sun, S.C., Brown, N.G., Xiong, Y., and Xu, Y.P. (2020). USP15 suppresses tumor immunity via deubiquitylation and inactivation of TET2. Sci. Adv. 6 1–13.

32. Huangfu, L., Zhu, H., Wang, G., Chen, J., Wang, Y., Fan, B., Wang, X., Yao, Q., Guo, T., Han, J., et al. (2024). The deubiquitinase USP15 drives malignant progression of gastric cancer through glucose metabolism remodeling. J. Exp. Clin. Cancer Res. 43, 235.

33. Cecchini, C., Pannilunghi, S., Tardy, S., and Scapozza, L. (2021). From Conception to Development: Investigating PROTACs Features for Improved Cell Permeability and Successful Protein Degradation. Front. Chem. 9, 672267.

34. Wiltshire, T.D., Lovejoy, C.A., Wang, T., Xia, F., O’Connor, M.J., and Cortez, D. (2010). Sensitivity to poly(ADP-ribose) polymerase (PARP) inhibition identifies ubiquitin-specific peptidase 11 (USP11) as a regulator of DNA double-strand break repair. J. Biol. Chem. 285, 14565–14571.

35. Ting, X., Xia, L., Yang, J., He, L., Si, W., Shang, Y., and Sun, L. (2019). USP11 acts as a histone deubiquitinase functioning in chromatin reorganization during DNA repair. Nuc. Acids Res. 47, 9721–9740.

36. Mah, L.J., El-Osta, A., and Karagiannis, T.C. (2010). gammaH2AX: a sensitive molecular marker of DNA damage and repair. Leukemia 24, 679–686.

37. Burgess, R.C., Burman, B., Kruhlak, M.J., and Misteli, T. (2014). Activation of DNA damage response signaling by condensed chromatin. Cell Rep. 9, 1703–1717.

38. Park, E.J., Chan, D.W., Park, J.H., Oettinger, M.A., and Kwon, J. (2003). DNA-PK is activated by nucleosomes and phosphorylates H2AX within the nucleosomes in an acetylation-dependent manner. Nuc. Acids Res. 31, 6819–6827.

39. Taccioli, G.E., Gottlieb, T.M., Blunt, T., Priestley, A., Demengeot, J., Mizuta, R., Lehmann, A.R., Alt, F.W., Jackson, S.P., and Jeggo, P.A. (1994). Ku80: Product of the XRCC5 Gene and Its Role in DNA Repair and V(D)J. Science 265, 1442–1445.

40. Maertens, G.N., El Messaoudi-Aubert, S., Elderkin, S., Hiom, K., and Peters, G. (2010). Ubiquitin-specific proteases 7 and 11 modulate Polycomb regulation of the INK4a tumour suppressor. Embo J. 29, 2553–2565.

41. Sowa, M.E., Bennett, E.J., Gygi, S.P., and Harper, J.W. (2009). Defining the human deubiquitinating enzyme interaction landscape. Cell 138, 389–403.

42. Inobe, T., Fishbain, S., Prakash, S., and Matouschek, A. (2011). Defining the geometry of the two-component proteasome degron. Nat. Chem. Biol. 7, 161–167.

43. Prakash, S., Tian, L., Ratliff, K.S., Lehotzky, R.E., and Matouschek, A. (2004). An unstructured initiation site is required for efficient proteasome-mediated degradation. Nature Struc. Mol. Biol. 11, 830–837.

44. Das, C., Hoang, Q.Q., Kreinbring, C.A., Luchansky, S.J., Meray, R.K., Ray, S.S., Lansbury, P.T., Ringe, D., and Petsko, G.A. (2006). Structural basis for conformational plasticityof the Parkinson’s disease-associatedubiquitin hydrolase UCH-L1. Proc. Natl. Acad. Sci. USA 103, 4675–4680.

45. Martinez-Fonts, K., Davis, C., Tomita, T., Elsasser, S., Nager, A.R., Shi, Y., Finley, D., and Matouschek, A. (2020). The proteasome 19S cap and its ubiquitin receptors provide a versatile recognition platform for substrates. Nat. Commun. 11, 477.

46. Beskow, A., Grimberg, K.B., Bott, L.C., Salomons, F.A., Dantuma, N.P., and Young, P. (2009). A conserved unfoldase activity for the p97 AAA-ATPase in proteasomal degradation. J. Mol. Biol. 394, 732–746.

47. Olszewski, M.M., Williams, C., Dong, K.C., and Martin, A. (2019). The Cdc48 unfoldase prepares well-folded protein substrates for degradation by the 26S proteasome. Commun. Biol 2, 29–37.

48. Li, H., Ji, Z., Paulo, J.A., Gygi, S.P., and Rapoport, T.A. (2024). Bidirectional substrate shuttling between the 26S proteasome and the Cdc48 ATPase promotes protein degradation. Mol. Cell 84, 1290–1303.

49. Kabuta, T., Furuta, A., Aoki, S., Furuta, K., and Wada, K. (2008). Aberrant interaction between Parkinson disease-associated mutant UCH-L1 and the lysosomal receptor for chaperone-mediated autophagy. J. Biol. Chem. 283, 23731–23738.

50. Zhang, L., Zhou, F., Drabsch, Y., Gao, R., Snaar-Jagalska, B.E., Mickanin, C., Huang, H., Sheppard, K.A., Porter, J.A., Lu, C.X., and ten Dijke, P. (2012). USP4 is regulated by AKT phosphorylation and directly deubiquitylates TGF-beta type I receptor. Nat. Cell Biol. 14, 717–726.

51. Virnau, P., Mirny, L.A., and Kardar, M. (2006). Intricate knots in proteins: Function and evolution. PLoS Comput. Biol. 2, e122.

52. Balzarini, M., Tong, J., Gui, W., Jayalath, I.M., Schell, B.B., and Kodadek, T. (2024). Recruitment to the Proteasome Is Necessary but Not Sufficient for Chemically Induced, Ubiquitin-Independent Degradation of Native Proteins. ACS Chem. Biol. 19, 2323–2335.

53. Qian, W.J., Liu, T., Monroe, M.M., Strittmatter, E.F., Jacobs, J.M., Kangas, L.J., Petritis, K., Camp, D.G., and Smith, R.D. (2005). Probability-Based Evaluation of Peptide and Protein Identifications from Tandem Mass Spectrometry and SEQUEST Analysis: The Human Proteome. J. of Proteome Res. 4, 53–62.

54. Keller, A., Nesvizhskii, A.I., Kolker, E., and Aebersold, R. (2002). Empirical Statistical Model To Estimate the Accuracy of Peptide Identifications Made byMS/MS and Database Search. Anal. Chem. 74 5383–5392.

